# Satiety Does Not Silence Food Attention: BMI Modulates Attentional Bias Toward High-Calorie Cues

**DOI:** 10.1101/2025.07.06.663349

**Authors:** Marc Ballestero-Arnau, Borja Rodríguez-Herreros, Toni Cunillera

**Author notes:** Corresponding autor: E-mail addresses (M. Ballestero-Arnau), (B. Rodríguez-Herreros), (T. Cunillera).

## Abstract

**Background:** Attentional bias (AB) for palatable foods has been linked to overeating, yet data on its modulation with body-mass index (BMI) and hunger/satiety are inconsistent. We tested whether hunger, sensory-specific satiety and BMI interact to shape automatic capture of attention by real high-calorie snack foods.

**Methods:** Two emotional-attentional-blink (EAB) experiments presented food distractors either 300 msec (lag-3) or 900 msec (lag-9) before a neutral target. Experiment 1 (N = 183; whole BMI range) randomly assigned participants to *taste* (non-satiated), *satiated* or *non-eating* conditions; the snacks eaten (or not eaten) subsequently re-appeared as distractors in the EAB task. Experiment 2 (N = 61; 31 overweight/obese, 30 normal-weight) manipulated hunger/satiety states and snack type (consumed vs. novel) orthogonally, in a double session design.

**Results:** Experiment 1— food images presented at lag-3 reduced target detection compared with lag-9 (OR = 0.61, CI 0.50–0.75, *p* <.01. Higher BMI predicted a larger AB when hungry, but a smaller AB when satiated (OR = 0.65, CI 0.48–0.88, p < 0.01). Experiment 2— For novel snacks, an interaction revealed that participants with overweight/obesity retained robust AB after satiation, whereas AB declined for participants with normal-weight (OR = 2.45, CI 1.09–5.51, p = 0.03). For snacks just eaten, AB remained significant in both groups (lag-3 vs. lag-9: OR = 0.37, CI 0.25–0.55, p <.001).

**Conclusions:** Real-food cues automatically biased attention regardless of metabolic state, with BMI modulating this effect: even when satiated, individuals with overweight/obesity continue to orient their attention toward both familiar and novel high-calorie foods. These findings suggest that satiety signals alone may be insufficient to curb attentional capture by food in obesity, highlighting the need for interventions that target attentional control and limit food availability.

## Introduction

The worldwide rise in obesity has intensified the search for cognitive mechanisms involved in the overconsumption of energy-dense foods despite balanced energy levels (1). One crucial mechanism is attentional bias (AB) — the tendency to allocate attention to motivationally salient cues while ignoring others (2). Recent research show that food-related AB can predict cravings, future energy intake and weight gain (3–5). However, there is mixed evidence regarding the link between body-mass index (BMI) and AB toward food cues, since some authors report individuals with overweight or obesity (OvOb) struggling to disengage from palatable food cues, leading to maladaptive behaviours (4,6), while others have found little or no group differences (7,8). A recent meta-analysis concluded that BMI explains only limited variance of AB (9).

One explanation for these inconsistencies could be that physiological states could modulate food-cue salience. For instance, hunger would increase the incentive value of calories (10) and would also amplify AB (11), consistent with evolutionary ideas that direct attention toward energy sources when resources are low (12,13). However, recent studies failed to replicate such effects (14,15), perhaps due to the high variability across paradigms (16). Equally important, satiety may down-regulate AB, but the existing evidence diverge: whereas in some studies satiated participants ignore food cues (17,18), in others, especially in OvOb samples, AB persists (19,20).

Another largely unexplored AB modulatory factor related to satiety is whether the satiety-related reductions of AB are stimulus-specific. Sensory-specific satiety (SSS) theory posits that the motivational value of food drops sharply once ingested, whereas uneaten alternatives remain appealing (21,22). In that sense, Di Pellegrino and colleagues observed lower interference from foods just eaten (18), but other study found that adults with obesity exhibited an AB even after consuming a nutritionally-matched liquid meal (19). Therefore, this issue still needs to be tested directly in individuals with overweight and obesity, who may continue to show strong AB by novel cues even after satiety (23,24).

To address these gaps, we employed a variant of the rapid-serial-visual-presentation (RSVP) task to induce an emotional attentional-blink (EAB) (25). During the RSVP, items appear in a single spatial location following a rapid succession; when an emotional distractor precedes a neutral target by ∼200-400 msec (lag-2/-3) the accuracy in detecting the appearance or absence of the target decay, and performance returns to baseline when the temporal distance between the distractor and target reaches ∼800-1000 msec (lag-8/-9) (26,27). Therefore, placing food distractors at lag-3 would be enough to indirectly assess attentional capture by food-distractor cues, whereas lag-9 could be used as baseline. The present study consisted of two experiments designed to shed light on the aforementioned inconsistencies, by manipulating hunger/satiety using a real-food approach with the aim to examine how BMI modulates food-driven AB.

### Experiment 1

To investigate the effects of hunger and satiation on attentional blink (AB), participants were assigned to one of three experimental conditions: *tast*e, *satiation*, or *non-eating*. In the *taste* condition, participants received a small portion of high-calorie snack foods, solely for taste appreciation and with the cover aim of eliciting a sustained feeling of hunger. In contrast, participants allocated to the s*atiation* condition were encouraged to eat snacks until feeling completely satiated. Finally, the *non-eating* group served as control condition in this experiment. Thus, participants shared the same table with other participants that were eating the snacks, and therefore they were indirectly exposed to food while performing an irrelevant task. In a subsequent phase of the experiment, images of the snacks previously consumed (or exposed) were presented as distractors during the RSVP task. In line with previous findings (28–31), we expected that all participants, regardless of the assigned condition, would show an AB effect (even in the non-eating condition) as it has been consistently observed that the mere exposure to others’ food consumption can also activate food-related cravings (32,33). Additionally, we hypothesized that participants in the *taste* condition would exhibit the strongest AB, since they would still be hungry. Finally, we hypothesized that BMI would modulate AB, with higher BMI associated with a more pronounced AB effect, particularly in the *taste* condition.

## Methods

### Participants

196 students from the Faculty of Psychology at the Universitat de Barcelona were recruited to participate in the study. Participants were excluded if they adhered to a vegan or vegetarian diet. Data from 13 participants were excluded because their accuracy on the experimental conditions without distractors was below 2.5 SD of the mean. Therefore, the final sample was composed of 183 participants covering the whole BMI range (see Supplementary Material) distributed across the *taste* (n = 49), *satiation* (n = 47) and non-eating (n = 87) conditions. Prior to the experiment, participants provided an informed consent approved by the local ethics committee. They were compensated with course credits for participant in the study.

### Stimuli and procedure

The stimuli employed in the experimental task comprised 19 colour images, extracted from the Blechert database (34), belonging to different image categories to form series in each trial, with or without the inclusion of the target stimulus (see Figure 1). Half of the series also included distractor images, consisting in pictures of real snack foods that participants previously consumed in the laboratory. There was a total of eight snacks, and their corresponding pictures grouped into two different sets of 4 snacks that were equally distributed across participants. Thus, the snacks from one set served as food in the pre-experimental conditions, while the pictures of those foods were used as distractor images in the RSVP task, and vice versa for the other set of snacks (see Supplementary Material).

**Figure 1.**
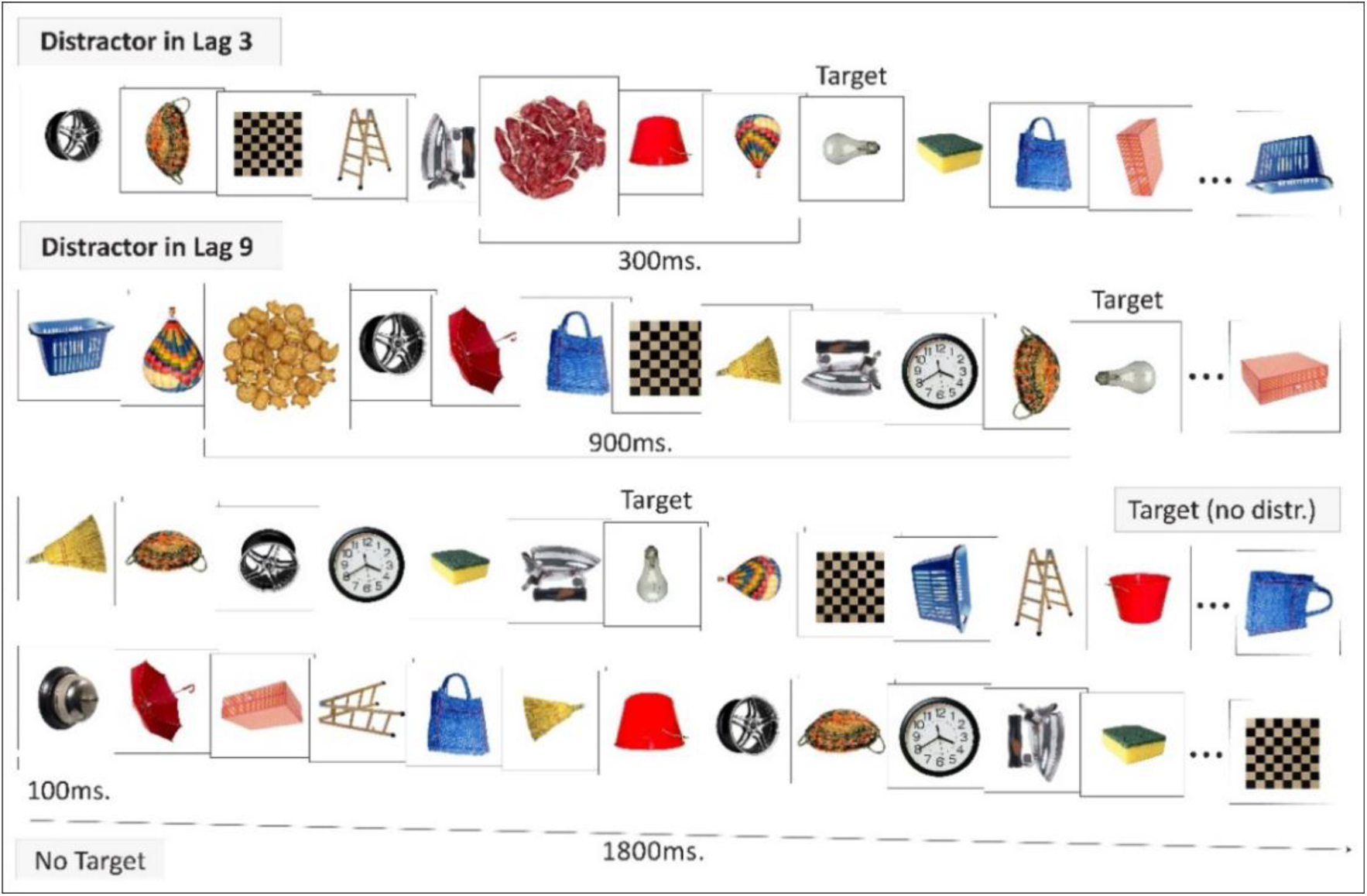
Example of each condition in the RSVP task created for the study. Series in the lag-3 condition: the distractor appeared 300 msec before the target (light bulb). Series in the lag-9 condition: the distractor was presented 900 msec before the target. Series in which the target was presented without the distractor. And finally, series in which neither the target nor the distractors were presented. Distractors could be either an image depicting an item from snack set 1 or snack set 2, depending on the assigned set. Although distractor images were size-matched to the other stimuli, they appear enlarged in the figure relative to their actual dimensions.

The participants were required to participate in the experiment following a fasting period of at least 4 hours (*M* = 6.9; SD = 3.6). Upon their arrival, all participants completed a survey providing their weight and height and indicating their subjective hunger level on a 10-cm line-scale, including intervals from 0 to 10 (0 indicating *no hunger at all*, and 10 being *very hungry*). Subsequently, half of the participants were randomly assigned to one of the two eating conditions (*taste* or *satiation*) and were then provided with one of two sets of snacks (see Figure S2A). Participants in the *taste* condition, received a small portion of snacks (∼73.9 kcal) to be consumed in approximately 5 minutes. Conversely, participants in the *satiation* condition were instructed to consume, within approximately 15 minutes, as much snack foods as they needed (up to ∼739.9 kcal) to feel completely satiated (see Figure S2B). Both, the experimental conditions and snack foods sets were counterbalanced across participants.

After eating all the snack foods or indicating they were satiated, participants were required to report on the snack’s *explicit liking* and *wanting* measures (35), using the same line-scale as in the previous measures (See supplementary Material). The other half of the sample, corresponding to those participants assigned to the *non-eating* condition, were requested to fill out the *Rational Experiential Inventory* (REI) questionnaire (36) at the time their partners were seated in front of them, eating at the same table. The questionnaire was used to keep participants busy while their peers were eating. Nevertheless, just before starting the experimental task, these participants were asked to indicate which foods were consumed by the peer sitting in front of them, and to report their *explicit wanting* about those snacks using the same line-scale that participants used in the *eating* condition. Participants in the non-eating condition differed in their indirect exposure time to the snack foods (*time-exposure*). Specifically, those participants paired with partners in the *taste* and *satiation* conditions were exposed for approximately 5 and 15 minutes, respectively. These *time-exposure* differences were considered and analysed to assess their potential impact on AB in the current study.

Upon finishing the pre-task requirements, all participants underwent the RSVP task. The experiment was conducted on an Intel i5 computer with Windows 10 Professional (64 bits) and using a routine programmed in PsychoPy (37). The experimental task was presented on a 24” LCD monitor with a refresh rate set at 60 Hz. The RSVP task consisted of a variant of the classical paradigm employed to elicit an EAB (14,15,31,38), in which participants must report the appearance or absence of a single target in every trial. Each series comprised 18 images displayed for 100 msec each, subtending on 10.8 degrees of visual angle. To mitigate potential habituation to the stimuli, images in the series were randomly and equitably rotated at 0°, 90°, 180°, and 270°. A total of 480 series were presented, with the target appearing in the position between the 6^th^ and 17^th^ images in 50% of the series.

A distractor image –the depicted a snack food– appeared within the series in 50% of trials when the target was presented. Thus, each of the four snack pictures were individually presented 30 times in 120 different trials, along with the other 17 neutral images in each series. The distractors were systematically presented three or nine positions before the target, i.e., lag-3 or lag-9, respectively (Figure 1). Brief breaks of approximately 1 minute were included after every 120 trials.

Finally, after completing the RSVP task, participants were asked again to rate their *subjective*-*hunger* level and to provide an estimation of the *calorie content* and the *consumption frequency* of all the food used in the experiment.

### Data analysis

Generalized Linear Mixed Models (GLMM) were computed separately to analyse participants’ task performance. All models incorporated always the condition *eaters* (taste vs. satiation) or *non-eaters* (short-exposure vs. long-exposure) and included the main *eaters* (reference: *satiation*), *lag* (*lag-3* vs. *lag-9*; reference: l*ag-9*) and BMI, as well as the interaction with the other two factors (See Supplementary Material).

## Results

An ANOVA including *snack foods* and task performance revealed a main effect of *lag* [*M lag-3* accuracy = 90.4% (SD = 8); *M lag-9* accuracy = 93.8% (SD = 5.3); *F*(1,180) = 43.85; *p* < 0.001; *η_p_^2^* = 0.20], with no effect of *snack foods* [*F*(1,180) = 0.28; *p* = 0.59; *η_p_ ^2^* = 0.001) or interaction [*F*(1,180) = 0.02; *p* = 0.88; *η_p_ ^2^*< 0.001]. Consequently, the two sets of snacks were collapsed for all subsequent analyses.

Next, the analysis for *hunger* revealed significant differences between *eaters* [taste *Δ*: *M* = –0.97, SD = 2.5; satiation *Δ*: *M* = –3.38, SD = 2.5; *t*(85) = 4.87; *p* < 0.001; *d* = 0.99], but not between *non-eaters* [*t*(94) = 1.79; *p* = 0.077; *d* = 0.38], indicating that hunger decreased only in satiated participants, thus validating the effectiveness of our experimental manipulation (Table 1). Finally, accuracy performing the general task without distractors was also assessed (M = 94.4%, SD = 4.8%) (See Supplementary Material).

**Table 1.**
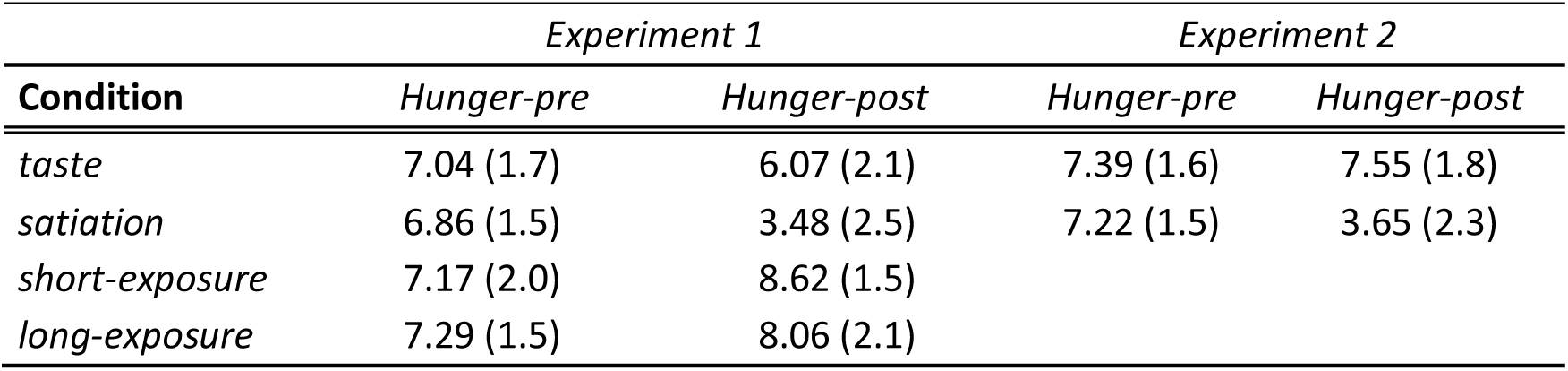
Mean values, standard deviations (in parentheses) in experiments 1 and 2 for pre-post-experiment measurements of hunger feeling.

### Eaters EAB performance (lag-3 vs. lag-9)

Results from the baseline model indicated that, relative to *lag-9*, performance at *lag-3* was predicted to be significantly lower (OR = 0.60, 95% CI [0.50, 0.73], *p* < 0.001) indicating an effective AB elicited by food distractors. The two *eaters’* conditions performed similarly (OR = 1.05, 95% CI [0.70, 1.57], *p* = 0.829), *lag* x *eaters* (OR = 0.98, 95% CI [0.74, 1.30], *p* = 0.897). The inclusion of BMI on the GLMM significantly improved the model fit (*χ*² = 9.96, *p* = 0.041) although BMI did not show a main effect (OR = 0.95, 95% CI [0.70, 1.29], *p* = 0.744), with task performance at *lag-3* remaining significantly lower (OR = 0.61, 95% CI [0.50, 0.75], *p* < 0.001). Moreover, the *lag* x *BMI* interaction was significant (OR = 1.28, 95% CI [1.02, 1.62], *p* = 0.037), as well as the interaction *lag x eating* x *BMI* (OR = 0.65, 95% CI [0.48, 0.88], *p* < 0.01). The predicted probabilities and log odds estimates (Table 2) indicated that, as BMI increased, the difference in task performance between *lag-3* and *lag-9* became larger in the *taste* condition. The reverse pattern was observed for *satiation*, suggesting that BMI mediated on the magnitude of the EAB effect as a function of hunger/satiety (Figure 2).

**Figure 2.**
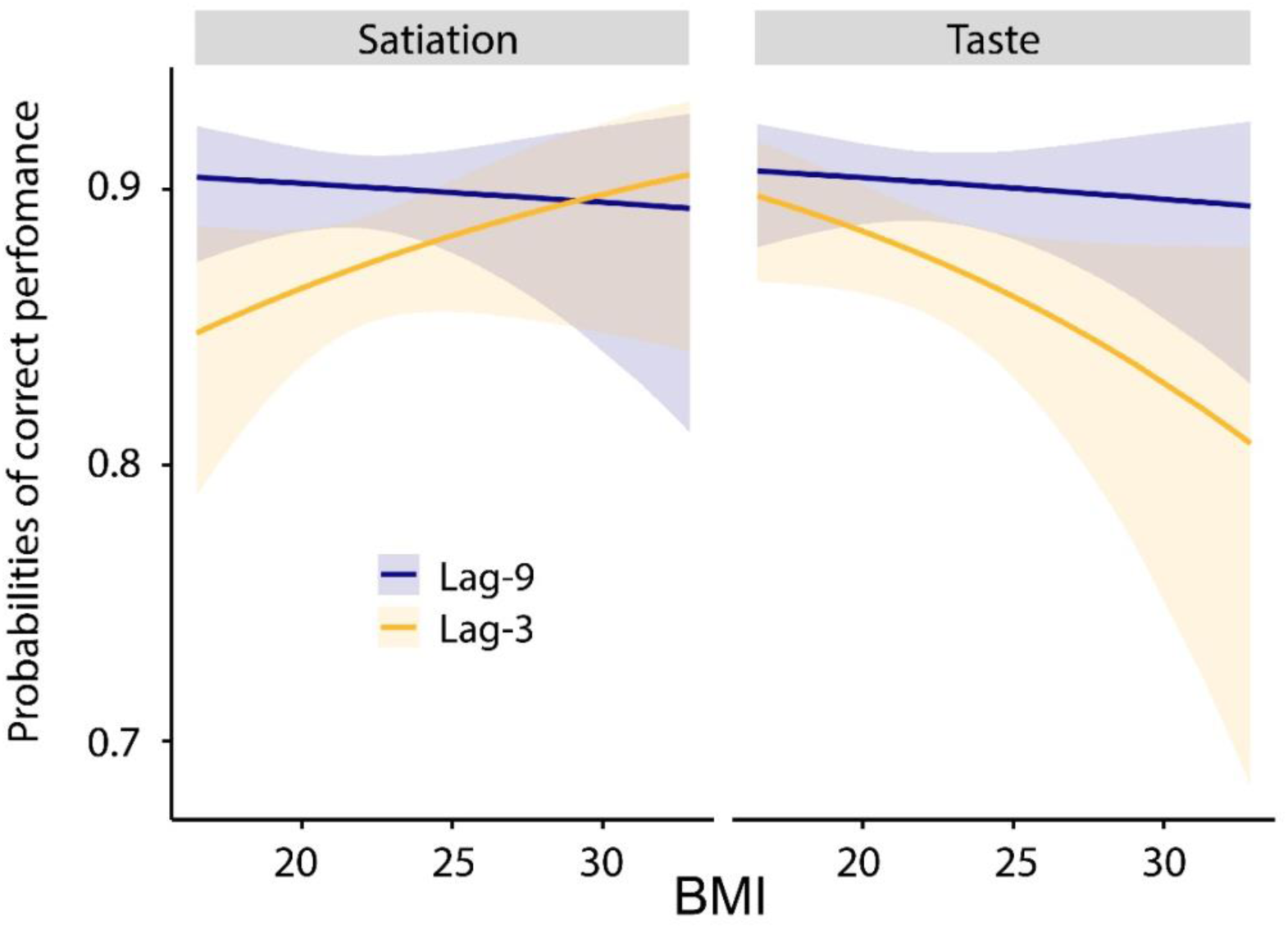
Experiment 1 predicted probabilities from the GLMM for correct responses with food distractors appearing in lag-3 and lag-9. Visualization of the three-way interaction *lag* x *eaters* x *BMI* represented separately by *eaters* (taste vs. satiation) conditions.

**Table 2.** Fixed effects estimates from the logistic mixed-effects model predicting attentional capture (lag-9 vs. lag-3) for eaters, including BMI. *Lag* reference level: *lag-9*. *Eaters* reference level: satiation. BMI is standardized.

| Predictor | Estimate (B) | SE | *z* | *p*-value |
| --- | --- | --- | --- | --- |
| Intercept | 2.99 | 0.15 | 20.26 | < .001 |
| **Lag** | -0.49 | 0.10 | -4.75 | < .001 |
| *Eaters* | 0.04 | 0.21 | 0.21 | .836 |
| *BMI* | -0.05 | 0.16 | -0.33 | .744 |
| *Lag* × *eaters* | -0.01 | 0.14 | -0.06 | .953 |
| *Lag* × *BMI* | 0.25 | 0.12 | 2.09 | .037 |
| *Eaters* × *BMI* | -0.01 | 0.21 | -0.04 | .966 |
| **Lag** × *eaters* × *BMI* | -0.43 | 0.15 | -2.80 | < .01 |

**Note.** Estimates are in log odds. Statistically significant effects are shown in bold.

### Non-Eaters general task performance and EAB

To assess the attentional effect of food distractor cues in *non-eaters*, we fitted a separated model. Results indicated that participants’ task performance was significantly lower in *lag-3* (OR = 0.66, 95% CI [0.54, 0.81], *p* < 0.001). No effects of *time*-*exposure* emerged (OR = 0.97, 95% CI [0.67, 1.43], *p* = 0.899; *lag* × *time*-*exposure*: OR = 0.97, 95% CI [0.73, 1.31], *p* = 0.869). Next, we evaluated an extended model for *non-eaters* including BMI and its interactions. Results indicated that incorporating BMI did not improve model fit (*χ*² = 1.22, *p* = 0.875).

## Discussion

Experiment 1 provided supporting evidence of food cues capturing participants attention, regardless of physiological states and BMI (8,28,31). Specifically, performance on the RVSP task declined when food distractors were presented shortly before the target, i.e. in the critical lag-3 condition, where the attentional blink is well-documented to occur (25,38,39). Furthermore, no differences were found between *taste* and *satiation* conditions until BMI was included in the GLMMs. Interestingly, higher BMI scores predicted a decrease in task performance in *lag-3* trials when hungry (*taste* condition). In contrast, the opposite pattern was found for satiated participants.

These findings align with persistent attentional capture from food-cues in participants with overweight and obesity (5,20) and are reminiscent of effects reported in substance-cue studies (3,40). However, even though our task provides an indirect measure of how satiation modulates AB, it remains uncertain whether the AB reduction is genuine and stimulus-specific, or an attenuation of responsiveness to any food cues (15,18), and whether BMI would moderate those effects (23,24).

### Experiment 2

We conducted a second experiment, now contrasting consumed and novel (non-consumed) high-calorie distractors. In a double session (*hunger* and *satiation*) within-subjects design, individuals with normal-weight (Nw) and overweight/obese (OvOb) completed the RSVP task. This design allowed us to test whether the previous encountered effect of satiation on the EAB was limited to familiar distractors or could otherwise be extended to novel stimuli, and whether such an effect would be modulated by BMI. It has been found that, under satiation, Nw individuals exhibit higher AB toward food than individuals with OvOb (19,41), which contrasts with the postulates of the SSS theory (15,18). Accordingly, we examined these two apparently opposite predictions: under a general AB account, participants with OvOb, once satiated on a given food, would continue to exhibit a strong EAB to novel and consumed foods, reflecting a broad attentional capture from food cues. In contrast, according to SSS theory, satiation would selectively reduce the AB toward foods just eaten while leaving novel foods unaffected, and this pattern should emerge irrespective of BMI if SSS operates independently (42).

## Methods

### Participants

70 students categorized as either Nw or OvOb were recruited. Participants with OvOb self-reported no diagnosis of diabetes, no history of mental health disorders or bariatric surgery, followed an omnivorous diet, and had a measured BMI greater than 27. Data from 9 participants were excluded from the selected sample because of exclusion criteria (see Experiment 1 methods). Therefore, the final sample was composed of 61 participants [31 OvOb (16 females), 30 Nw (25 females); *M* age = 22.7 years (SD = 3.6); range: 18–34 years; *t*(59) = 1.51; *p* = 0.131; *d* = 0.19] (see Supplementary material for details). All participants were required to attend the two experimental sessions under counterbalanced *taste* and *satiation eating* conditions. Participants’ subjective hunger was measured in each session before and after completing the experiment.

### Stimuli and procedure

The same two sets of snacks from Experiment 1 were used for consumption. However, the distractors presented in the RSVP task corresponded to either foods consumed during that session (half of trials), or foods from another set not consumed in any of the two experimental sessions. Thus, two new sets of four stimuli were extracted from the Blechert database (34). Set 1 consisted of: burger, pasta, cookie, and chocolate bar; Set 2: Hot dog, pizza, pralines, and ice cream. All sets of distractors were matched in both physical and nutritional composition properties (see Supplementary Material). In each session, the distractors presented in the RSVP task corresponded to one of the two consumed food sets, while the other half were novel foods from one of the two new sets. The assignment of food sets and *eating* conditions was counterbalanced across participants and experimental sessions.

The RSVP task remained identical to Experiment 1, except for the addition of the new factor *distractor-type* (specific vs. novel). To equally present distractors from the two types in all possible positions, a total of 384 series were presented with the eight snack foods appearing 12 times each, in 96 different trials. Participants first completed the assigned *eating* condition (taste/satiation), and immediately afterwards they performed the RSVP task. Same fasting period as in experiment 1 was implemented (*M* = 6.9; SD = 3.6).

## Results

Comparison of consumed snacks revealed only a main effect of *lag* [*F*(1,59) = 34.70; *p* < 0.001; *η_p_*^2^ = 0.37], not influenced by *snack type* [*F*(1,59) = 0.91; *p* = 0.344; *η*_p_ ^2^ = 0.02; *lag* x *snack type*: *F*(1,59) = 0.31; *p* = 0.581; *η*_p_ ^2^ = 0.005]. The same comparison was conducted for the non-eaten snacks, yielding again a main effect of *lag* [*F*(1,59) = 17.19; *p* < 0.001; *η*_p_ ^2^ = 0.23], with neither *snack type* influence [*F*(1,59) = 1.34; *p* = 0.251; *η*_p_ ^2^ = 0.02], nor *lag* x *snack type interaction*[*F*(1,59) = 0.02; *p* = 0.882; *η_p_*^2^ <.001]. Therefore, the two novel sets were collapsed for all the analyses.

In terms of hunger/satiety, comparisons in *hunger difference* revealed a main effect of *eating* [*M taste Δ* = 0.15 (SD = 1.95); *M satiation Δ* =-3.57 (SD = 2.8); *F*(1,59) = 98.49; *p* <.001; *η_p_*^2^ = 0.63], but no effect of *group* [*F*(1,59) = 0.69; *p* = 0.411; *η_p_*^2^ = 0.01], *eating* x *group* [*F*(1,59) = 0.47; *p* = 0.495; *η_p_*^2^ <.001]; highlighting the successful hunger/satiety manipulation (see Table 1).

### Novel food distractors and EAB

For novel distractors, a GLMM was fitted to predict correct task performance, as a function of *lag* (*lag-3* vs. *lag-9*; reference: *lag-9*), *group* (OvOb vs. Nw; reference: Nw), and *eating* (taste vs. satiation; reference: *satiation*), including always participant ID as a random factor (see Table 3). Results showed no effects of *lag* (OR = 0.95, 95% CI [0.62, 1.46], *p* = 0.820), *group* (OR = 0.69, 95% CI [0.39, 1.22], *p* = 0.199), or *eating* (OR = 1.53, 95% CI [0.95, 2.48], *p* = 0.082). There was, however, a significant *lag* x *group* (OR = 0.49, 95% CI [0.28, 0.85], *p* = 0.011) and *lag* x *eating* interaction (OR = 0.51, 95% CI [0.27, 0.96], *p* = 0.037), as well as a three-way interaction *lag* x *group* x *eating* (OR = 2.45, 95% CI [1.09, 5.51], *p* = 0.03), suggesting that the influence of *group* on the effect of *lag* depended on the hunger/satiety state. The inspection of predicted probabilities indicated that participants with OvOb would show higher AB than Nw, as predicted by a decrease in the odds of correct responses in *lag-3* trials, when presented with novel distractors during the *satiation* condition (Figure 3; Table 3).

**Figure 3.**
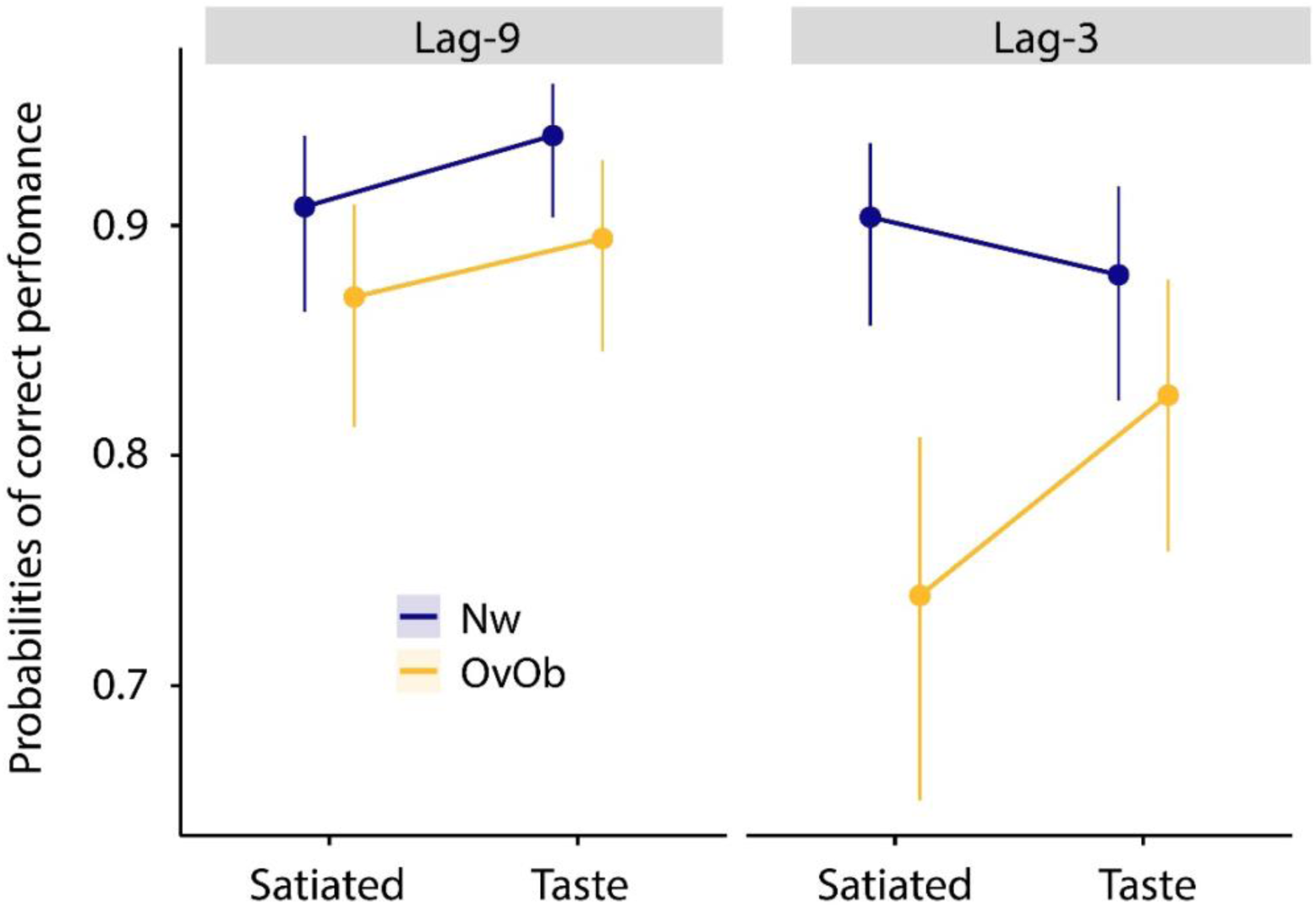
Experiment 2 predicted probabilities from the GLMM for correct responses with novel distractors and for *lag-3* and *lag-9*. Visualization of the three-way interaction *lag* x *group* x *eating* represented separately by *eating* conditions and *lag*.

**Table 3.** Fixed-effects estimates (Log Odds) from the GLMM analysing *task* performance for *lag* incorporating novel distractors in the model. *Lag* reference level: *lag-9*. *Group* reference level: Nw. *Eating* reference level: satiation.

| Predictor | Estimate (B) | SE | z | p-value |
| --- | --- | --- | --- | --- |
| (Intercept) | 3.034 | 0.219 | 13.87 | < .001 |
| *Lag* | -0.050 | 0.220 | -0.23 | 0.820 |
| *Group* | -0.377 | 0.293 | -0.71 | 0.199 |
| *Eating* | 0.427 | 0.246 | 1.74 | 0.082 |
| ***Lag* × *group*** | -0.710 | 0.280 | -2.54 | **0.011** |
| ***Lag* × *eating*** | -0.672 | 0.322 | -2.09 | **0.037** |
| *group* × *eating* | -0.198 | 0.125 | -1.19 | 0.534 |
| ***Lag* × *group* × *eating*** | 0.898 | 0.413 | 2.17 | **0.030** |

**Note.** Estimates are in log odds. The factors reaching Statistically significant effects are shown in bold.

### Specific eaten food distractors and EAB

Results from the GLMM for eaten distractors showed a main effect of *lag* (OR = 0.37, 95% CI [0.25, 0.55], *p* < 0.001), indicating that participants in both groups had significantly lower odds of giving correct responses in *lag-3* trials. The effect of *group* was also significant (OR = 0.44, 95% CI [0.24, 0.82], *p* < 0.01), with participants in the OvOb group showing lowered odds of correct task performance. In contrast, no effects of *eating* (OR = 1.13, 95% CI [0.70, 1.81], *p* = 0.621) were found, and none of the interaction terms reached statistical significance (All *p-values >.15*). Overall, these results indicate that the attentional capture from the images representing the foods just eaten was more pronounced in participants with overweight/obesity in both hunger and satiated states, with all participants showing a decrease in the odds of providing correct responses when specific food distractors appeared in *lag-3* (see Supplementary Material, Table S2).

## Discussion

Findings from Experiment 2 show that satiation did not consistently attenuate the AB to high-calorie foods in the OvOb group. Notably, participants with OvOb still exhibited a strong AB toward novel (non-consumed) snacks when satiated, whereas this bias subsided in participants with Nw. These results suggest that, in participants with OvOb, attentional capture by food cues might generalize to any high-calorie cue and persist despite physiological fullness, echoing prior evidence of enduring cue-driven responsivity in obesity (6,16) and theories of heightened reactivity to palatable food stimuli (43,44).

### General Discussion

We investigated how hunger, satiation, and BMI modulated the AB toward high-calorie food cues. In Experiment 1, all participants displayed strong AB toward food distractor cues, particularly under hungry conditions. This bias was amplified in individuals with higher BMI and was reduced after eating, suggesting some modulation of food-related attention by the current metabolic state and BMI. Notably, even participants in the non-eating condition—who only observed others consuming snacks—showed elevated but weaker AB. These findings highlight that both internal (e.g., hunger) and external (e.g., social cues) influences can trigger AB toward food stimuli (5,20,32,33).

Experiment 2 expanded on these findings by addressing a central question: is the observed drop in AB specific to the snacks just consumed, or does it extend to other high-calorie foods? To answer this, participants completed a double session experiment (hungry and satiated), which allowed us to compare the AB for previously consumed versus novel food distractors in Nw and OvOb targeted groups. Results revealed a divergence between groups: whereas Nw participants showed an overall reduction in AB when satiated, OvOb individuals maintained a strong AB—particularly for non-consumed (novel) snack items. These findings suggest that satiation alone may not fully attenuate food attentional capture in individuals with higher BMI which is consistent with prior work showing that individuals with OvOb exhibit persistent AB to palatable foods, even when satiated (16,19,41). At the same time, our results challenge the classical SSS theory, which posits that once a particular food is consumed, its attentional and motivational value diminishes more than uneaten alternatives (15,18), even regardless of inter-individual differences (42). In our study, AB remained present even for the foods just eaten, particularly in the OvOb group, suggesting that consumption does not fully suppress attentional prioritization of those cues.

One plausible explanation for the persistence of AB under satiety could be that the mere exposure to real cues may have primed attention, potentially overriding satiation (45,46). In that sense, theories of goal activation propose that appetitive cues— independently of hunger—can activate a mindset associated with seeking behaviour (46–48). This could also partly explain why AB persisted for consumed stimuli and even intensified for novel ones in the OvOb group, unlike Nw (19,49,50). Although we cannot confirm this mechanism with our current design, it offers a plausible interpretation that could be further tested. Moreover, because the RSVP task began immediately after eating, the abovementioned cue-priming may have dominated while satiety signals were still emerging (51,52). In that sense, brief exposure to palatable foods is suggested to capture attention for several minutes, whereas markers of satiety typically peak 15– 30 min after a meal (10,53).

## Conclusions

The current study reveals that hunger, satiation, BMI and stimulus novelty interactively modulate food-related AB, unveiling a key finding: even when satiated, people with overweight/obesity are still attracted to high-calorie foods (35,43,53)—particularly those not yet consumed. Targeting this cue-driven attentional capture through interventions—such as attentional bias modification training (17,40) and environmental measures to limit high-calorie food availability—should be central to strategies addressing maladaptive eating behaviours in the so-called obesogenic environment we live in (1).

## ACKNOWLEDGEMENTS

This research was supported by research grants from the Spanish Government (Ministerio de Ciencia e Innovación) to T.C. (grant number: PID2021-127743OB-I00).

## Supplementary material

### Experiment 1

we describe here the sample of the experiment in terms of statistical power and BMI. We also provide further information on the stimuli and task procedure, as well as the amount of food consumed to assume participants satiation. Results from target-detection baseline performance (GLMM testing effects of target presence/absence) are also reported here, and follow-up tests of relevant subjective measures (hunger, explicit wanting, explicit liking, perceived calories, consumption frequency) to verify that none of these variables altered the primary results.

## METHODS

### Participants

G*Power analysis indicated that a sample of 181 participants was sufficient to assess the effect of 2 within and 1 between condition, and their interaction terms (with a power = 95% and an a-priori alpha set at *p* = 0.05), for a moderate effect size estimate (*η*_p_ ^2^ = 0.06). The distributions of *BMI* scores were similar across experimental conditions [*M* BMI = 21.91 (SD = 3.2); range: 16.49 – 34.45; *F* (3,178) = 0.52; *p* = 0.667; *η_p_^2^*= 0.008].

### Stimuli and Procedure

The four stimuli in each set of snack foods shared identical nutritional composition in terms of Kcal/100gr. [*M* set 1 = 461.5 (SD = 124.17); *M* set 2 = 397.75 (SD = 221.48); *t*(6) = 0.5; *p* = 0.633; *d* = 0.36], Carbohydrates/100gr. [*M* set 1 = 33.66 (SD = 34.5); *M* set 2 = 33.42 (SD = 34.61); *t*(6) = 0.01; *p* = 0.994; *d* = 0.01], Fat/100gr. [*M* set 1 = 28.66 (SD = 19.85); *M* set 2 = 22.12 (SD = 24.33); *t*(6) = 0.42; *p* = 0.692; *d* = 0.29], and Protein/100gr.

[*M* set 1 = 14.87 (SD = 9.99); *M* set 2 = 13.8 (SD = 7.78); *t(*6) = 0.17; *p* = 0.871; *d* = 0.12], with its counterpart, as stated in the FBBD website (https://fddb.info). The physical properties of those food images (size, brightness, contrast, complexity, and spatial frequency) were quantified using OpenCV (1) and Scikit-image (2) Python libraries. One-way ANOVAs were conducted for each parameter across all sets of images (Neutral, snack foods set-1 and snack foods set-2) revealed non-statistical difference for any parameter: size [*F(*2,24) = 0.89; *p* = 0.424; *η_p_* ^2^ = 0.07]; brightness [*F(*2,24) = 0.39; *p* = 0.685; *η_p_*^2^ = 0.03]; contrast [*F(*2,24) = 1.89; *p* = 0.174; *η*_p_ ^2^ = 0.14]; complexity [*F(*2,24) = 0.69; *p* = 0.51; *η_p_* ^2^ = 0.05]; spatial frequency [*F(*2,24) = 0.54; *p* = 0.589; *η_p_* ^2^ = 0.04].

Apart from hunger levels, the rest of subjective measures were taken using the same 10-cm line scale. In the case of *Liking*: “how much do you liked the eaten snack foods” (0 meant *not liking at all*, and 10 signified *delicious*); for *wanting*: “how much more of those snacks foods you desire to eat at that moment” (0 indicated they would *not eat any more of this food*, and 10 signified a strong desire to continue eating *more of these snack foods*); for *perceived calories*: “How much kcal. do you think each of the eaten snacks had” (with 0 representing *low-calorie content* and 10 denoting *high-calorie content*); and finally, for *consumption frequency* a 5-point Likert scale was used (*Rarely, once a month, once a week, several times per week, Daily*).

### Food consumed (participants satiation)

In the *taste* condition, only 12 participants completed the consumption of the entire snack food servings, while the remaining participants left uneaten food on their plates [in grams: *M* = 26.04; SD = 24.7]. In terms of kcal consumed, participants had eaten ∼657 kcal (SD = 51.17).

## RESULTS

### Eaters general task performance (target presence or absence)

Results from the GLMM in baseline task performance showed that *target* presence significantly increased the odds of correct responses (OR = 1.22, 95% CI [1.06, 1.40], *p* < 0.01). However, *eating* did not significantly impact participants’ task performance (OR = 1.21, 95% CI [0.90, 1.63], *p* = 0.214), *target* x *eaters* (OR = 0.95, 95% CI [0.78, 1.17], *p* = 0.642; Table S1), indicating that the advantage for target-present trials in comparison to target-absent trials was similar whether participants had eaten or not.

### Non-eaters general task performance (target presence or absence)

Results also revealed that *target-presence* significantly increased the odds of correct responses (OR = 1.62, 95% CI [1.39, 1.89], *p* < 0.001). In contrast, the effect of *time*-*exposure* was not found to be significant (OR = 1.04, 95% CI [0.75, 1.44], *p* = 0.798); *target-presence* x *exposure*: OR = 0.84, 95% CI [0.68, 1.04], *p* = 0.104). These results indicate that the advantage for target-present trials was equivalent for both time exposures conditions.

### Eaters GLMMs to further explore subjective measures

To control for possible confounds, *t-tests* for the subjective measurements obtained during the experiment (*hunger, explicit wanting, explicit liking, perceived calories, and consumption frequency*) were computed separately between *eaters* and *non-eaters*. Only those measures reaching statistical significance were included in the GLMMs to examine whether they could improve model fit and therefore, indicate an influence on the obtained results.

Results revealed, apart from *hunger*, significant differences in *consumption frequency* [*taste*: *M* = 2.24, SD = 0.44; *satiation*: *M* = 2.01, SD = 0.48; *t*(94) = 2.41, *p* = 0.018].

However, likelihood-ratio tests indicated that neither *hunger* (*χ*² = 5.24, *p* = 0.732) nor *consumption frequency* (*χ*² = 9.83, *p* = 0.277) improved the final model’ fit.

### Non-eaters GLMMs and subjective measures

We found that neither *hunger*, *wanting*, *consumption frequency*, nor *perceived calories* differed significantly between *non-eaters’* conditions (all *p*-*values* > 0.07). Consequently, no further models were tested.

#### Experiment 2

We further describe here the composition of the two groups (normo-weight vs. overweight/obese), BMI distributions, as well as statistical power; detailed information about stimuli and task procedure (including food and non-food images nutritional and visual properties comparisons); and the amount of snack consumed in both groups and conditions. Results from the GLMM of target-detection performance are also detailed, as well as a summarized table with averaged values between groups and conditions in the two *lags* tested (lag-3 vs. lag-9).

## METHODS

### Participants

G*Power calculations indicated that 52 participants were sufficient to analyze 3 within-subjects and one between-factor with an identical power, a-priori alpha set, and effect size estimate as in Experiment 1.

BMI scores were different between groups [*M* Nw = 21.88 (SD = 1.81) range: 18.05 – 24.65; *M* OvOb = 31.61 (SD = 4.29) range: 27.07 – 42.44; *t*(59) =-11.32; *p* < 0.001; *d* = - 2.89].

### Stimuli properties and procedure

No statistical difference for nutritional composition and physical properties were found between the different sets of food images. In terms of physical properties, no differences were found between sets (including non-food stimuli) for *size* [*F(*3,27) = 0.86; *p* = 0.473; *η_p_*^2^ = 0.09], *brightness* [*F(*3,27) = 0.58; *p* = 0.637; *η*_p_ ^2^ = 0.06], *contrast* [*F(*3,27) = 1.36; *p* = 0.278; *η*_p_ ^2^ = 0.13], *complexity* [*F(*3,27) = 0.79; *p* = 0.51; *η*_p_ ^2^ = 0.08], or *spatial frequency* [*F(*3,27) = 0.83; *p* = 0.48; *η*_p_ ^2^ = 0.08]. Regarding nutritional composition, food sets were matched in *Total Kcal.* [*F(*3,12) = 1.10; *p* = 0.386; *η*_p_ ^2^ = 0.22], *protein content* [*F(*3,12) = 0.85; *p* = 0.492; *η_p_* ^2^ = 0.18], *fat content* [*F(*3,12) = 0.44; *p* = 0.731; *η_p_* ^2^ = 0.10], and *carbohydrates content* [*F(*3,12) = 0.33; *p* = 0.804; *ηp*^2^ = 0.08].

### Food consumed (participants satiation)

Only 6 and 12 participants from the Nw and OvOb conditions, respectively, consumed all the snacks served in the *satiation* condition. The remaining participants left uneaten food on their plates [Nw *M* = 33.96 gr; SD = 22.03; OvOb *M* = 41.11 gr; SD = 22.05; *t*(41) =-1.04; *p* = 0.303; *d* =-0.32]. In kcal. [Nw *M* = ∼659 kcal.; SD = 50.3; OvOb *M* = ∼643 kcal.; SD = 50.8; *t*(41) = 0.44; *p* = 0.66; *d* =-0.15].

## RESULTS

### General task performance (target presence vs. absence)

A GLMM was built to test baseline *task performance* considering BMI as a categorical predictor (group: *Nw* vs. *OvOb*). Results revealed that *target presence* or *absence* did not affect the odds of correct responses (OR = 1.10, 95% CI [0.88, 1.37], *p* = 0.406). Importantly, the main effect of *group* was non-significant (OR = 0.85, 95% CI [0.55, 1.33], *p* = 0.479), indicating that task accuracy in trials without distractor was similar between groups. However, in the *taste* condition participants were more likely to respond correctly on target-present trials than on target-absent trials (OR = 1.22, 95% CI [1.02, 1.47], *p* = 0.032), indicating that hunger reduced target-detection errors. None of the two or three-way interactions were statistically significant (all *p-values* >.19).

### GLMMs and subjective measures

No statistically significant differences were found between groups in neither *hunger, explicit wanting, explicit liking, perceived calories, and consumption frequency* (all *p*-*values* > 0.09). Therefore, no further models were tested.

---

**Figure S1.**
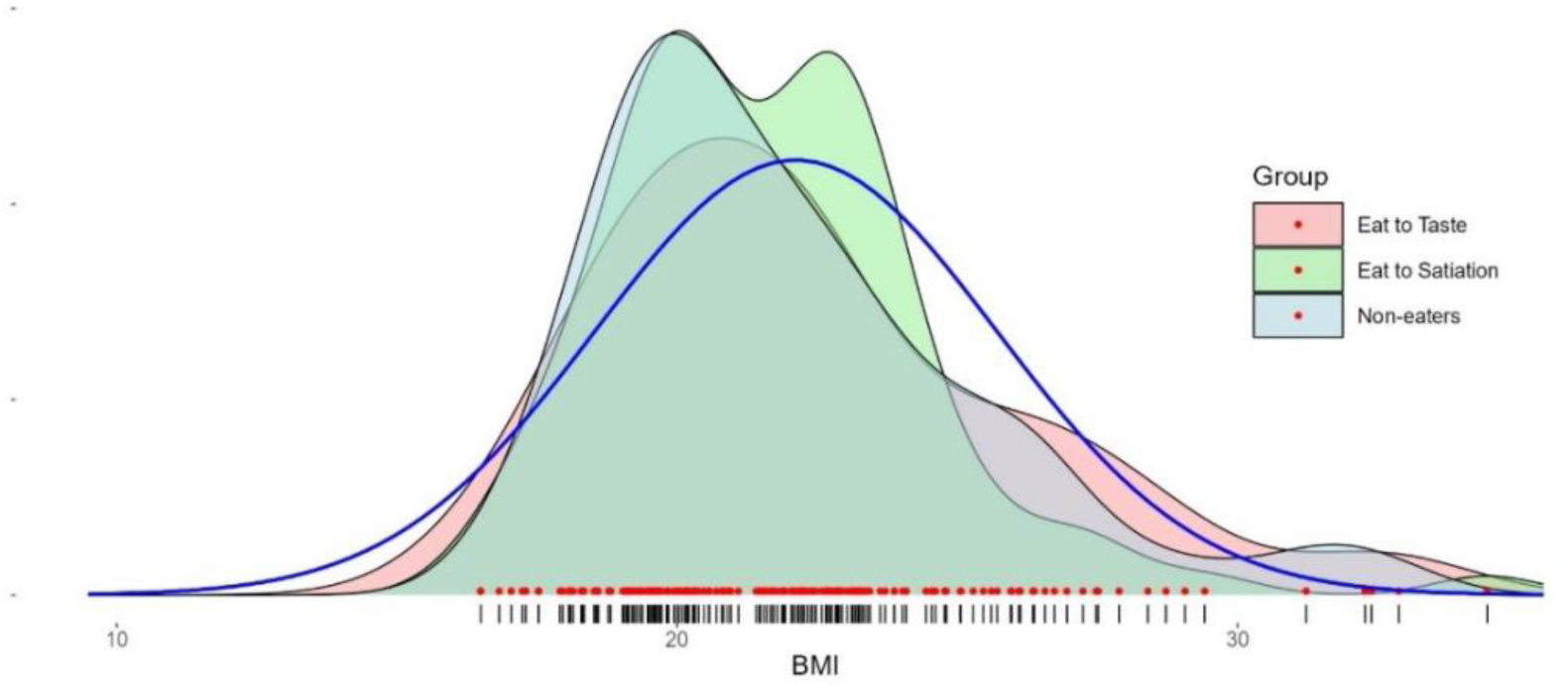
Equal BMI distributions for the three main conditions in Experiment 1. The blue curve depicts the theoretical Gaussian generated form the overall BMI mean and standard deviation pooled across conditions. Data from the two non-eating conditions (short-and long-exposure) are collapsed.

**Figure S2.**
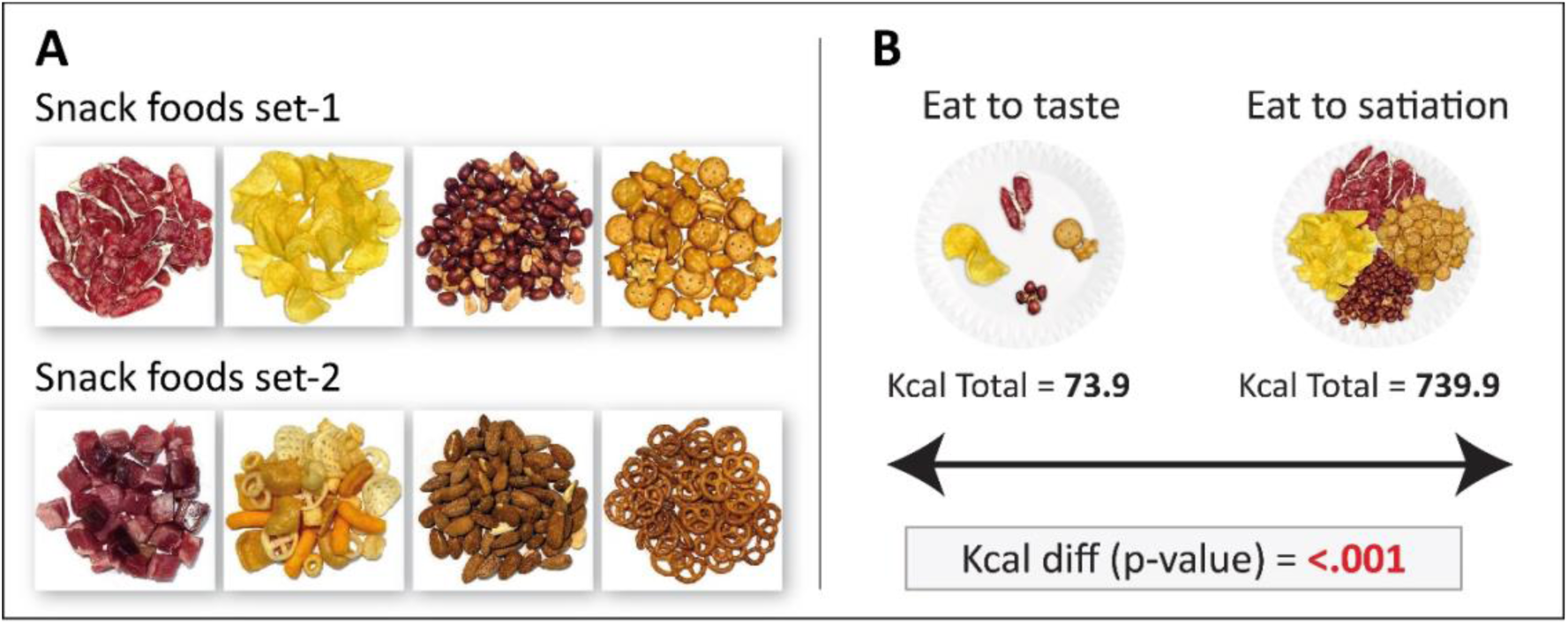
*A*) Snack-food Set 1 used as distractor images (top row): dry-cured sausage, potato’s chips, peanuts, and mixed savory crackers. Snack-food Set 2 (bottom row) comprised cured ham, corn snacks, almonds, and uniform savory crackers. *B*) Illustration of the portions provided in the *taste* (left) and *satiation* (right) conditions, showing their respective energy content (73.9 kcal. Vs. 739.9 kcal). The caloric difference between portions was significant [*t(*14) =-7.44; *p* < 0.001; *d* =-3.72].

**Fig. S3.**
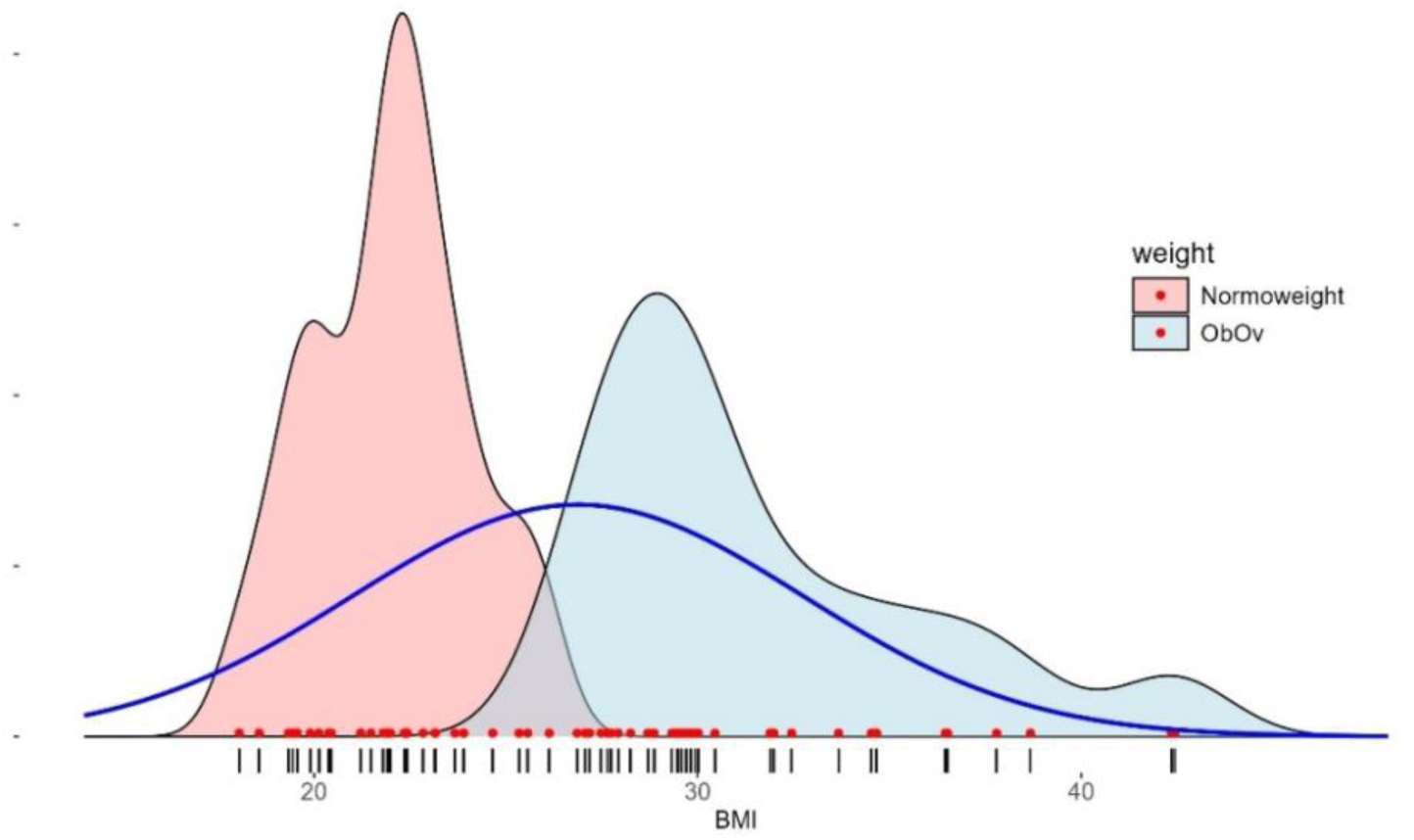
BMI distributions for Nw and OvOb in Experiment 2. The blue line shows the theoretical distribution of BMI values assuming they follow a Gaussian distribution based on the overall mean and standard deviation from the two groups.

**Table S1.** Fixed effects estimates from the logistic mixed-effects model predicting task performance for *eaters*.

| Predictor | Estimate (B) | SE | z | p-value |
| --- | --- | --- | --- | --- |
| *Intercept* | 2.87 | 0.11 | 26.52 | < .001 |
| ***Target presence*** | 0.20 | 0.07 | 2.81 | **.005** |
| *Eaters* | 0.19 | 0.15 | 1.24 | .214 |
| *Target presence × Eaters* | 0.05 | 0.10 | 0.47 | .642 |

**Note.** Target absence is the reference of the model. *Eaters* reference level: *satiation*. Estimates are in log odds. Factors reaching statistical significance are shown in bold.

**Table S2.** Averaged performance (%) for *group, eating* conditions (satiation vs. hunger/taste, and *snack type* (novel vs. specific) in Experiment 2. Standard deviations are shown in parentheses.

| Group | Eating | Snack Type | Lag-3 | Lag-9 |
| --- | --- | --- | --- | --- |
| Normo-weight | Satiation | Novel | 93.75 (5.7) | 94.03 (6.7) |
|  |  | Specific | 87.36 (10.6) | 94.58 (6.8) |
|  | Taste | Novel | 92.22 (11.3) | 95.97 (4.9) |
|  |  | Specific | 87.78 (14.7) | 95.14 (5.6) |
| Overweight / Obesity | Satiation | Novel | 84.82 (15.6) | 91.94 (10.2) |
|  |  | Specific | 83.2(15.9) | 89.78 (11.3) |
|  | Taste | Novel | 89.52 (11.4) | 93.41 (7.6) |
|  |  | Specific | 84.95 (15.0) | 92.47 (7,7) |

